# A deep learning and co-conservation framework enable discovery of non-canonical Cas proteins

**DOI:** 10.1101/2025.09.30.679098

**Authors:** Beibei He, Chen Qi, Yuanyuan Feng, Dong Liu, Fangrong Wu, Zongan Wang, Dan Wang, Zhen Yue, Yong Zhang, Hongxia Lan, Yue Zheng, Yuxiang Li

## Abstract

CRISPR–Cas systems are central to prokaryotic adaptive immunity, widely harnessed for biotechnology. Yet, their vast and uncharacterized diversity, especially non-canonical variants, impedes full exploitation. Here we present BioPrinCRISPR, a class-agnostic computational framework leveraging gene co-conservation, protein domain co-occurrence, and embedding similarity to identify and characterize CRISPR–Cas systems across prokaryotic genomes. Applying BioPrinCRISPR to over one million bacterial genomes, we uncovered extensive canonical and uncharacterized systems, revealing a rich landscape of atypical Cas proteins and novel domain architectures. Notably, we identified recurrent fusion proteins with unique enzymatic combinations, suggesting roles in regulatory control or nucleic acid remodeling. Experimental validation of two divergent Cas13an-like effectors demonstrated RNA knockdown capacity in human cells, confirming our framework’s predictive power. These findings expand the functional repertoire of CRISPR-associated proteins and highlight unexplored modes of microbial immunity. BioPrinCRISPR thus stands as a powerful tool for comprehensively mapping CRISPR–Cas diversity, offering new insights into prokaryotic defense and facilitating discovery of novel candidates for next-generation genome engineering. An accompanying interactive web platform was also developed to facilitate data exploration.

## Introduction

Clustered Regularly Interspaced Short Palindromic Repeat (CRISPR) and genes encoding CRISPR-associated (Cas) proteins was first identified to be an immune system in bacteria^[1]^. Decades of research have revealed that CRISPR-Cas systems utilize RNA molecules transcribed from the CRISPR array to guide Cas protein in cleaving foreign DNA or RNA ^[2]^. This gRNA-guided DNA/RNA cleavage ability is not only central to the CRISPR-Cas immune mechanism but also a key feature that has been harnessed to develop genome-editing tools with unparalleled precision ^[3]^. This unique defense mechanism was then reprogrammed to an efficient and site-specific genome editing tool, now one of the most widely used technologies in diverse fields, including gene therapy, agriculture and fundamental biological research^[4-7]^. With 2 classes encompassing 6 types of CRISPR-Cas systems identified so far^[2]^, and several new subtypes being characterized each year^[4, 8, 9]^, CRISPR-Cas continues to evolve as a powerful tool for genome editing.

The accumulation of characterized CRISPR-Cas systems has profoundly advanced the development of bioinformatic algorithms for their annotation from the rapidly growing genomic sequencing data. Methods like Basic Local Alignment Search Tool (BLAST) and Hidden Markov Models (HMMs) have been foundational, enabling large-scale identification of known Cas protein homologs and their associated loci ^[10-12]^. A wealth of useful HMM-based tools has been developed in the recent years, including CasFinder ^[13]^, CRISPRCasFinder ^[14]^ and CRISPRCasTyper ^[15]^. More recently, predicting Cas protein using protein language model (PLM) has emerged as an powerful approach ^[16, 17]^, leveraging pretrained large protein language models such as ESM2 ^[18]^ and ProtBert ^[19]^ to capture complex sequence features. However, while these reference sequence-based methods have paved the way for extensive CRISPR-Cas annotation, their performance is inherently dependent on the quality and diversity of the known sequences used for building HMM profiles or training sets. Consequently, these approaches face significant challenges in identifying highly divergent or non-canonical systems that do not conform to predefined models. Their reliance on sequence or structural homology inherently limits the ability to detect systems with unique domain architectures, novel fusion proteins, or unconventional functional mechanisms. Furthermore, the rapid expansion of genomic data continually outpaces the manual updating of reference databases, leading to an increasing number of uncharacterized Cas proteins that remain elusive to conventional annotation workflows.

Here, we revisited the fundamental principle of CRISPR-Cas system in defending viral DNA and proposed a computational framework to define CRISPR-Cas system, accompanied by a domain-orientated Cas protein prioritization method, named BioPrinCRISPR (bio-principle-informed CRISPR-Cas system discovery). BioPrinCRISPR annotates CRISPR-Cas system in a class-independent manner, allowing for the discovery of novel Cas proteins and systems beyond classic CRISPR-Cas systems. We demonstrated the ability of BioPrinCRISPR by analyzing 1,067,859 bacterial assemblies from NCBI database. The results showed that the sequence space of classic Class 2 Cas proteins is largely unexplored, including type II, type V and type VI. We also revealed large number of new systems that do not fit into the current CRISPR-Cas category. BioPrinCRISPR and the dataset we introduced would contribute to the discovery of new CRISPR-Cas systems and expanding the genome editing toolbox.

## Results

### Co-occurrence and co-conservation between CRISPR array and Cas protein define CRISPR-Cas system

We hypothesized that the functional association between CRISPR arrays and Cas proteins would manifest as conserved co-occurrence patterns across diverse genome. CRISPR immune systems involve intensive Cas protein-RNA interactions, based on which we proposed that similar Cas proteins are associated with similar CRISPR arrays. A typical CRISPR-Cas system consists of a CRISPR array which features varying number of repeats and spacers and co-occurred proteins, including effector module and adaptation module ^[20]^. In adaptation stage, Cas1 and Cas2 complex is essential for the cleavage of viral DNA and the generation of new spacers ^[21]^. In the CRISPR RNA (crRNA) biogenesis stage, the pre-crRNA transcribed from the CRISPR array is processed to generate mature crRNA ^[22]^. In Class I systems, this processing is typically mediated by Cas6 or similar endoribonucleases ^[23-25]^, whereas in Class II systems, the effector proteins themselves (including Cas9, Cas12 and Cas13) possess instrinsic RNA-processing capabilities or rely on accessory proteins for crRNA maturation ^[26-30]^. This maturation process facilitates the formation of effector-RNA complexes that are crucial for targeting and cleaving foreign genetic materials. In the whole process, comparability between each entity is the foundation for spacer generation, crRNA and Cas-RNA complex formations. We hypothesized that such comparability is realized via co-conservation between CRISPR array and the associated Cas proteins, i.e., for similar Cas proteins, their array sequences are also similar, defined as co-conservation.

To validate our hypothesis, we analyzed 769 Cas9 family CRISPR systems, centering the CRISPR locus on the array and extending it 10 kb upstream and downstream, thereby covering both Cas and non-Cas proteins. We then conducted sequence similarity analysis (SSN) of both arrays and all the proteins from these CRISPR systems (**Figure 1A**). We define co-conservation (c) as the proportion of common elements between array clusters and protein clusters relative to the total number of elements in the protein cluster. A high c value indicates a strong association between a given Cas protein and its CRISPR array. For 769 known Cas9 systems, the c values for Cas9, Cas2 and Cas1 are 0.83, 0.71 and 1.0, respectively, indicating a high co-conservation between them (**Figure 1B**). Of note, the high c value of Cas1 is in agreement with the fact that Cas1 is one of the key proteins in generating spacer and processing the array-transcribed pre-crRNA in adaptation and expression stages. This result validated that similar arrays are associated with similar Cas proteins.

**Fig. 1.**
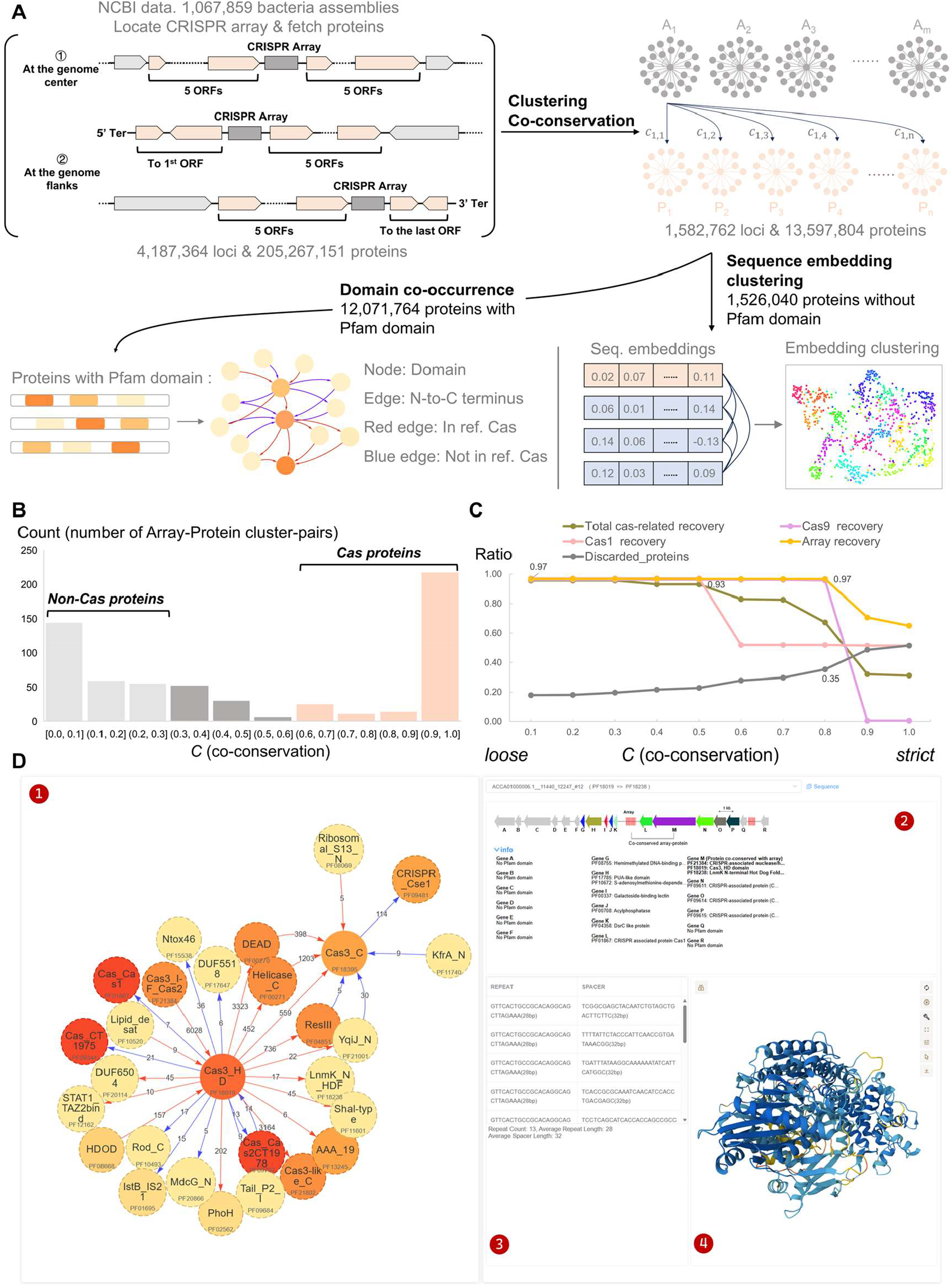
Co-conservation analysis of CRISPR arrays and Cas proteins in CRISPR-Cas9 systems. **(A)** Schematic of the workflow to assess co-conservation between CRISPR arrays (blue rectangle) and ±5 ORFs flanking the array are included (red arrows). Sequence similarity networks and Pfam domain clustering were used to define protein clusters. **(B)** Histogram showing the distribution of co-conservation C values between arrays and proteins across 769 Cas9 systems, highlighting peaks near 0 and 1. **(C)** Recall of Cas9, Cas1 and CRISPR arrays at varying co-conservation thresholds. **(D)** Interactive subnetwork page of the BioPrinCRISPR web application, integrating (1) an overview of the protein domain co-occurrence subnetwork centered on the selected domain, (2) gene organization of the operon containing the representative protein, (3) co-conserved CRISPR array information associated with the operon, and (4) ESMfold-predicted three-dimensional structure of the representative protein.

We noticed that the distribution of all-to-all c values exhibited a bimodal pattern, where the data is concentrated at two distinct ends, near 1 and near 0 (**Figure 1B**). This result suggests that the array-protein clusters tend to cluster around these two values, indicating two distinct behaviors, i.e., the proteins around a set of similar arrays are either similar (c nears 1) or divergent (c nears 0). This phenomenon aligns with the biological principle of the CRISPR-Cas system, where the proteins surrounding the array are either functionally associated with the immune process or unrelated to it. Co-conservation value is thus can be used to distinguish array-related proteins from array-non-related ones.

We evaluated the recall of Cas9, Cas1, and the CRISPR locus from 769 known Cas9 systems across varying thresholds of c, ranging from 0.1 to 1.0. As c increased, the recall of the array remains constant at 0.97 until c reaches 0.8. A sharp decrease of Cas1 recall was found at c equals to 0.5, where the total Cas recall also undergoes a slightly decrease to 0.93. The c value of 0.5 was thus selected to keep a high recall of both Cas proteins and a decent discarded protein ratio (**Figure 1C**).

### BioPrinCRISPR annotates bacterial CRISPR-Cas system

Here, we developed BioPrinCRISPR, a comprehensive computational workflow designed to identify and characterize canonical and non-canonical CRISPR-Cas systems in a class-independent manner by leveraging co-conservation and domain co-occurrence patterns. This approach builds upon the principle that conserved co-occurrence between CRISPR arrays and neighboring proteins can serve as a robust indicator for CRISPR system annotation, independent of known classifications. Using this strategy, we analyzed 1,067,859 bacterial assemblies from the NCBI database. CRISPR arrays were first detected using MinCED ^[31]^, and the flanking proteins were extracted to form array–protein sequence pairs.. A co-conservation analysis was performed to filter for consistently co-conserved pairs, resulting in 13,597,804 proteins and 1,582,762 loci.

To further investigate functional associations, we performed a domain co-occurrence analysis across the retained loci. Protein domains were annotated using HMMER, and overlapping or redundant domain hits were resolved through position-based and statistical filtering to ensure high-confidence domain assignments. Domain co-occurrence metrics were then calculated, quantifying the frequency and conservation of domain pairs within CRISPR-associated genomic regions. These metrics were used to construct a domain-level co-occurrence network, in which nodes represent protein domains and edges denote statistically supported co-presence. Known Cas domains were highlighted as references to indicate the discovery status of adjacent domains. Notably, a subset of proteins (1,526,040) lacked identifiable Pfam domains, prompting the use of sequence embedding-based similarity network analysis to explore remote homology and novel domain structures. Candidate systems can be prioritized based on novel domain composition, distant homology to known Cas proteins, compact size, or atypical organizational features. BioPrinCRISPR thus provides a scalable and unbiased framework for expanding the current landscape of CRISPR-Cas system discovery.

We noticed that Class one CRISPR-Cas system, including Cas1, Cas3 and Cas5, dominates the overall distribution of the entire domains. Hallmark domains of Class two CRISPR-Cas system, the HNH endonuclease and RuvC endonuclease and other CRISPR associated proteins ranked at the top. These data suggest that BioPrinCRISPR is able to capture CRISPR-Cas system independent of reference sequences. In addition to known CRISPR-related proteins, a wealth of non-canonical proteins is also abundantly found in the loci, including elongation factor, radical SAM superfamily, 4Fe-4S domain, glycosyl transferase and others. The functions of these enzymes are not directly associated with classic CRISPR-Cas system in the immune system in bacteria. However, the co-occurrence and co-conservation of these enzymes with array may imply biological importance.

Leveraging the BioPrinCRISPR framework, we established an interactive web application (URL: https://crispr-384817688195.asia-east2.run.app), deployed on Google Cloud Run, that comprehensively visualizes the protein domain co-occurrence network derived from the NCBI database, allowing users to perform advanced filtering, querying, and data retrieval. To mitigate redundancy, proteins were initially grouped based on their domain annotation profiles and subjected to a two-tier clustering strategy within each group: a primary clustering at 90% sequence identity to remove highly similar duplicates, followed by a secondary clustering at 30% sequence identity to delineate potential subfamilies. Representative proteins were selected as the top five largest cluster centroids for each domain combination. The application enables users to select domains of interest as central nodes and access their associated secondary subnetworks (Fig. 1D), which present detailed information including subnetwork topology, operon structures of representative proteins, linked CRISPR array characteristics, and predicted three-dimensional protein structures generated using ESMfold^[18]^. This platform offers a robust and scalable resource for systematic exploration of CRISPR-Cas-associated protein domain relationships, advancing the discovery and functional characterization of system diversity and novelty.

### Domain co-occurrence revealed novel fusion patterns in Cas proteins

Having established the utility of co-conservation for identifying CRISPR-associated proteins, we then sought to dissect their functional architecture through domain co-occurrence analysis, which provides insights into novel protein fusions and functional modules. Many Cas proteins are multidomain enzymes, a structural complexity that underpins their diverse functions in bacterial defense, such as DNA interaction, unwinding, and cleavage. To explore the domain co-occurrence of identified proteins, we used Cas proteins from CasPedia ^[32]^ (1,498 sequences) and CasPDB ^[33]^ (15,051 sequences) as known reference to to evaluate the novelty of co-occurring domains found in the domain co-occurrence network. Domain co-occurrence network analysis of candidates from the NCBI database revealed numerous domain fusions were absent from the CasPedia and CasPDB (**Figure 2**). In Class 2 system, HNH endonuclease domain plays a pivotal role in DNA cleavage. In the domain co-occurrence network, we noticed that the HNH-helicase_C domain fusion has not been previously observed in the CasPedia or CasDB databases. Further analyze the corresponding system from *Butyricicoccus pullicaecorum* DSM 23266 revealed that Cas1 (BpuE), Cas2 (BpuD) and Cas7 (BpuG) were encoded in the same operon. Notably, the domain architecture of BpuC features a DEAD helicase at N-terminus, a helicase_C domain in the middle and a HNH domain at C-terminus (**Figure 2A, 2B**). Such unique domain orchestration may imply an alternative mechanism in defending bacteriophages. We also found that HNH-Cas5 and HNH-Cys1 fusion patterns are also absent from the reference data. A previous research has demonstrated that such fusions are functional in DNA cleavage ^[34, 35]^.

**Fig. 2.**
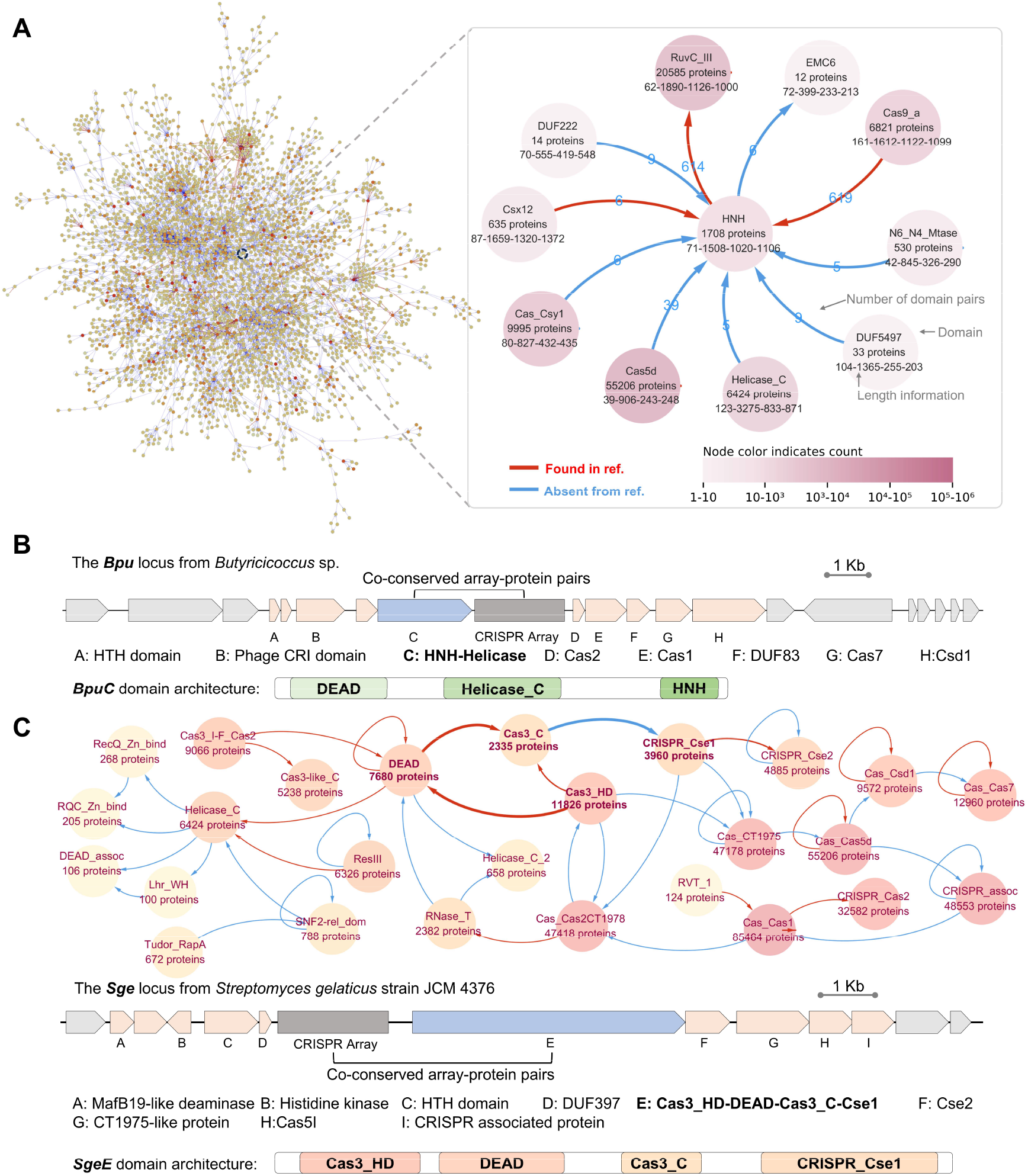
BioPrinCRISPR reveals non-canonical CRISPR-Cas systems via co-conservation and domain co-occurrence. **(A)** Domain co-occurrence network constructed from Pfam-A annotated protein domains. Known Cas domains (e.g., Cas1, Cas3, HNH, RuvC) and known domain co-occurrence edges are highlighted as red. **(B)** Domain architecture of the BpuC protein showing fused DEAD helicase, Helicase-C, and HNH nuclease domains, and its corresponding genomic locus. **(C)** Domain architecture of the SgeE protein, composed of fused Cas3_HD, DEAD helicase, Cas3_C, and Cse1 domains, and its corresponding genomic locus. The domain co-occurrence edges of SgeE are highlighted in bold. The network depicted here is a simplified overview; the full detailed network is available in the Supplementary Information.

Class1 system forms multi-subunit effector complexes to mediate target recognition and interference. However, in our analysis, we observed a remarkable deviation from the canonical multi-subunit configuration. Specifically, we noted that some of these discrete proteins, which are typically part of larger complexes, can fuse into a single protein unit, leading to new functional combinations not previously characterized. Examples of such fusions include the Cas_HD-Cas2 and Cas3_C-Cse1 proteins, which combine domains that are usually present in distinct effector proteins, as well as Csd1-Cas7 and several other novel domain co-occurrences (**Figure 2C**). These fused proteins, which represent a class of non-canonical or hybrid Cas proteins, may indicate the presence of more versatile or specialized systems that could have evolved to optimize CRISPR interference in specific microbial environments. The emergence of such domain fusions suggests a level of functional diversity within Class 1 CRISPR systems that has yet to be fully explored and may point to the existence of unique interference mechanisms tailored to specific ecological or evolutionary pressures.

### Domain Co-occurrence Network Illuminates Cas-Related Functional Modules

To systematically explore non-canonical CRISPR-like systems, we constructed a domain co-occurrence network, in which each node represents a protein domain and each edge indicates the co-occurrence of two domains within the same protein or operon. The resulting network contained X nodes and Y edges, forming 371 communities identified by the Louvain algorithm ^[36]^ (modularity = 0.942). The modularity score reflects the extent of community structure, calculated as:

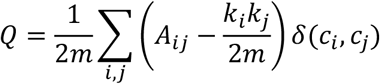

where *A*_*ij*_ is the weight of the edge between node *i* and node *j, k*_*i*_ and *k*_*j*_ are their degrees, m is the total number of edges, and *δ*(*c*_*i*_, c_*j*_) = 1 if nodes *i* and *j* belong to the same community, 0 otherwise. Higher modularity reflects stronger-than-random intra-community connectivity.

Among the 371 communities, 40 contained at least one known Cas domain, indicating a modular organization associated with CRISPR functionality (**Figure 3A**). Within these Cas-enriched communities, we identified 537 non-Cas domains co-localized with Cas components. Functional annotation of these domains revealed enrichment for nucleic acid-binding motifs (e.g., HTH_3, RRM_1) and domains of unknown function (e.g., DUF442, DUF4900), suggesting potential roles in regulation, effector targeting, or structural assembly.

**Fig. 3.**
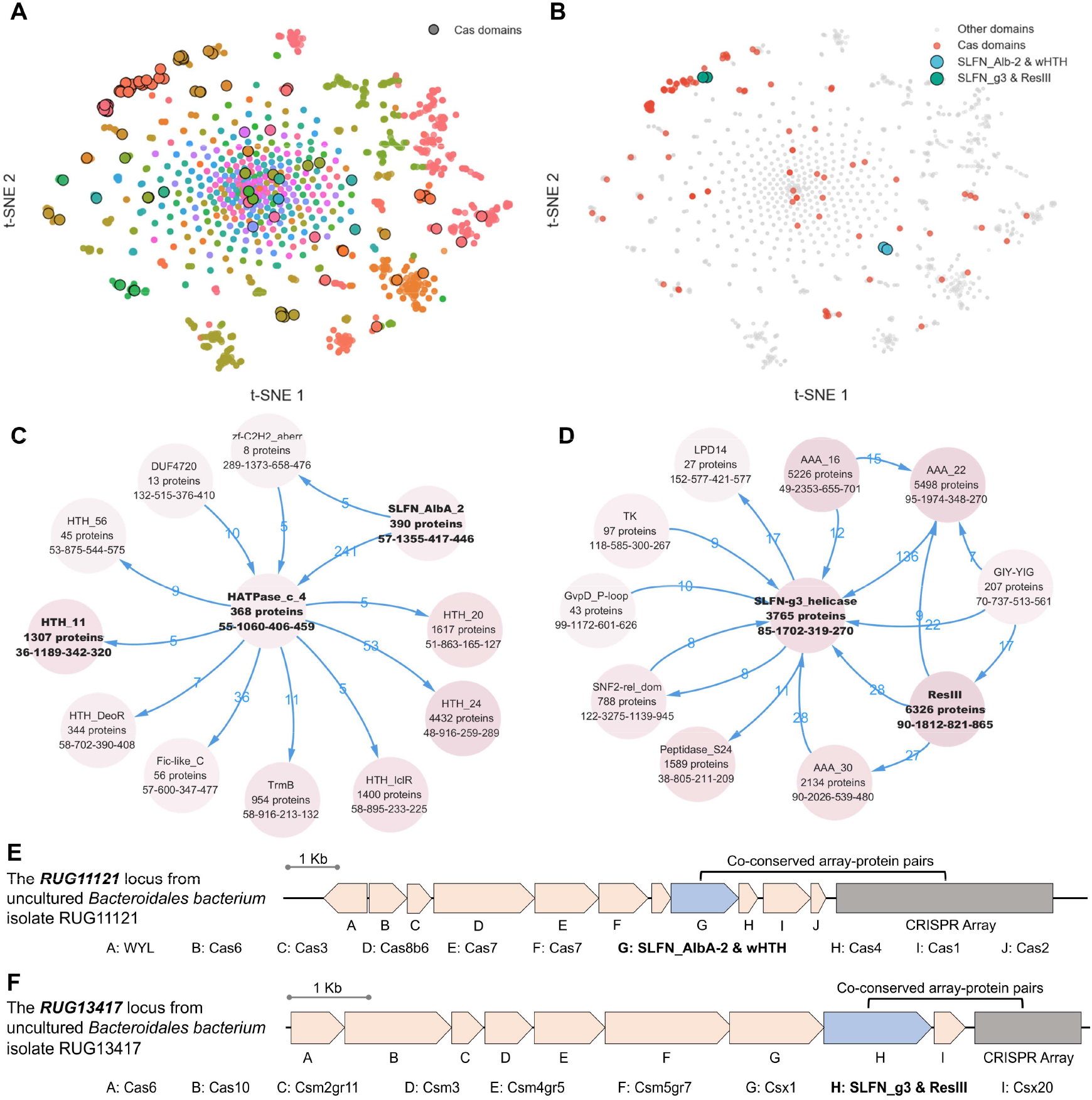
Domain co-occurrence network and representative Schlafen-like proteins in CRISPR loci. **(A)** t-SNE visualization of Louvain community partitioning of the domain co-occurrence network. **(B)** Relative positions of two Schlafen-like protein classes in the Louvain communities shown by t-SNE. **(C)** Detailed subnetwork of co-occurring proteins containing SLFN_AlbA-2 and wHTH domains. Node names shown in **bold** indicate proteins associated with these specific domains. **(D)** Detailed subnetwork of co-occurring proteins containing SLFN-g3_Helicase and ResIII domains. Node names shown in bold indicate proteins associated with these specific domains. **(E)** Representative genomic loci examples of SLFN_AlbA-2 and wHTH domain co-occurrence protein. **(F)** Representative genomic loci examples of SLFN-g3_Helicase and ResIII domain co-occurrence protein.

To prioritize candidate domains, we employed both topological and representation learning-based strategies. Centrality metrics—such as degree and betweenness—highlighted network hubs that may function as architectural scaffolds. In parallel, we analyzed domain fusion and adjacency patterns, identifying novel domain combinations absent from current CRISPR-Cas annotations.

We further trained Node2Vec ^[37]^ embeddings on the domain co-occurrence graph and introduced a Cas-likeness score to quantify the proximity of each domain to known Cas components in the latent embedding space. For a domain x, its Cas-likeness is defined as:

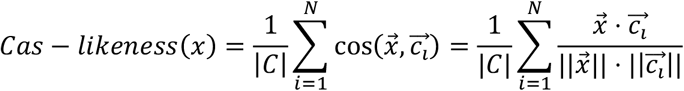

Where 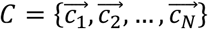 is a curated set of embedding vectors corresponding to experimentally validated Cas domains. High Cas-likeness scores indicate strong embedding similarity to canonical Cas proteins. Dimensionality reduction via t-SNE (perplexity = 30) on the 64-dimensional embeddings confirmed that high-scoring domains co-localize with known Cas modules in latent space, supporting their functional relevance.

Guided by a multi-layered prioritization framework, we systematically applied BioPrinCRISPR to protein domain-encoding loci across complete microbial genomes. This comprehensive survey revealed recurrent, non-canonical domain architectures that frequently co-localize with CRISPR arrays and core Cas gene modules, pointing to potential functional integration within diverse CRISPR-Cas systems.

Unexpectedly, within these prokaryotic genomes, we identified a subset of proteins annotated with Schlafen (SLFN)-related domains—despite the Schlafen family traditionally considered vertebrate-specific (**Figure 3B**). In eukaryotes, Schlafen proteins share high sequence homology but diverse functions and differential expression across tissues and species. The Schlafen family is known to participate in a broad range of biological processes, including DNA replication, cell proliferation, immune and interferon responses, viral restriction, and sensitivity to DNA-targeting anticancer agents ^[38]^.

A particularly notable finding was the co-occurrence of a Schlafen_AlbA-2 domain (PF04326, hereafter referred to as the AlbA-2 domain) and a winged helix-turn-helix (wHTH) motif (PF13412) within the same protein (Figure 3D). This domain pair was repeatedly identified within operons adjacent to Type I CRISPR-Cas gene clusters. The AlbA-2 domain, characterized by conserved glutamic acid residues, is predicted to have RNase activity, suggesting a role in RNA cleavage or processing ^[39]^. The accompanying wHTH motif, a canonical nucleic acid-binding fold, likely contributes to sequence-specific recognition of nucleic acid targets ^[40]^. Their co-localization, together with consistently high Cas-likeness scores, supports a dual functional role in target recognition and regulatory modulation. Interestingly, AlbA-2–wHTH-containing operons were often found at Class I CRISPR-Cas loci, where the core gene set—including Cas5, Cas7, and Cas8—remained intact. In these contexts, the AlbA2 fusion protein consistently appeared in syntenic positions typically associated with Class I accessory subunits. This pattern suggests that the AlbA-2–wHTH fusion may act as an auxiliary enhancer, augmenting the functionality of canonical Class I systems. Genomic neighborhood analysis across diverse taxa revealed conserved spatial associations with CRISPR repeats and core effector genes, suggesting this fusion protein constitutes a previously uncharacterized accessory factor within a Class I effector module. Given its predicted RNase activity and nucleic acid-binding motif, AlbA-2–wHTH-containing operons potentially bridge RNA cleavage, target detection and nucleic acid processing or transcriptional regulation.

In another set of loci, we identified another recurrent fusion protein that combines a Schlafen-like DNA/RNA helicase domain (PF09848) with a Type III restriction enzyme Res subunit-like domain (PF04851) (**Figure 3E**). This chimeric architecture was consistently observed within Type III CRISPR-Cas operons and, in a subset of cases, co-localized with core interference components from Class I systems, such as Cas3, Cas5, or Cas10. The Schlafen-like helicase domain, reminiscent of those found in eukaryotic immune regulators, likely mediates nucleic acid unwinding and contributes to the resolution of replication-associated stress ^[41-43]^. In parallel, the Res subunit domain, typically linked to ATP-dependent DNA translocation and endonucleolytic activity, implies functional coupling of helicase and nuclease activities ^[44, 45]^. Their co-occurrence suggests a modular effector that integrates substrate remodeling with targeted interference, potentially augmenting canonical Class I functions. The conserved domain architecture and its repeated proximity to Cas operons across diverse microbial genomes point to a previously uncharacterized accessory module with relevance to the evolution and diversification of CRISPR-based immunity.

### Characterization of Unannotated Proteins Through Semantic Embedding Clustering

To expand the discovery of CRISPR-Cas effectors beyond domain-annotated proteins, we leveraged a two-step computational pipeline integrating language model-based embeddings and unsupervised clustering. Protein sequences without Pfam-A domain annotations were first embedded using the ESM-2 protein language model, which captures sequence-level biochemical and structural features. The resulting fixed-length embeddings, derived by averaging the final hidden states of the model, were then clustered alongside a curated reference set of Class 2 Cas proteins (Types II, V, VI).

We applied k-means clustering to partition the embedding space and used UMAP for dimensionality reduction and visualization of the protein relationships. Proteins exhibiting high similarity to Cas proteins (i.e., low Euclidean distances) were prioritized as candidates for novel Cas-like functions. Interestingly, several unannotated proteins clustered with known Cas12 and Cas13 variants, suggesting shared functional features despite the absence of detectable Pfam domains (**Figure 4A**).

**Fig. 4.**
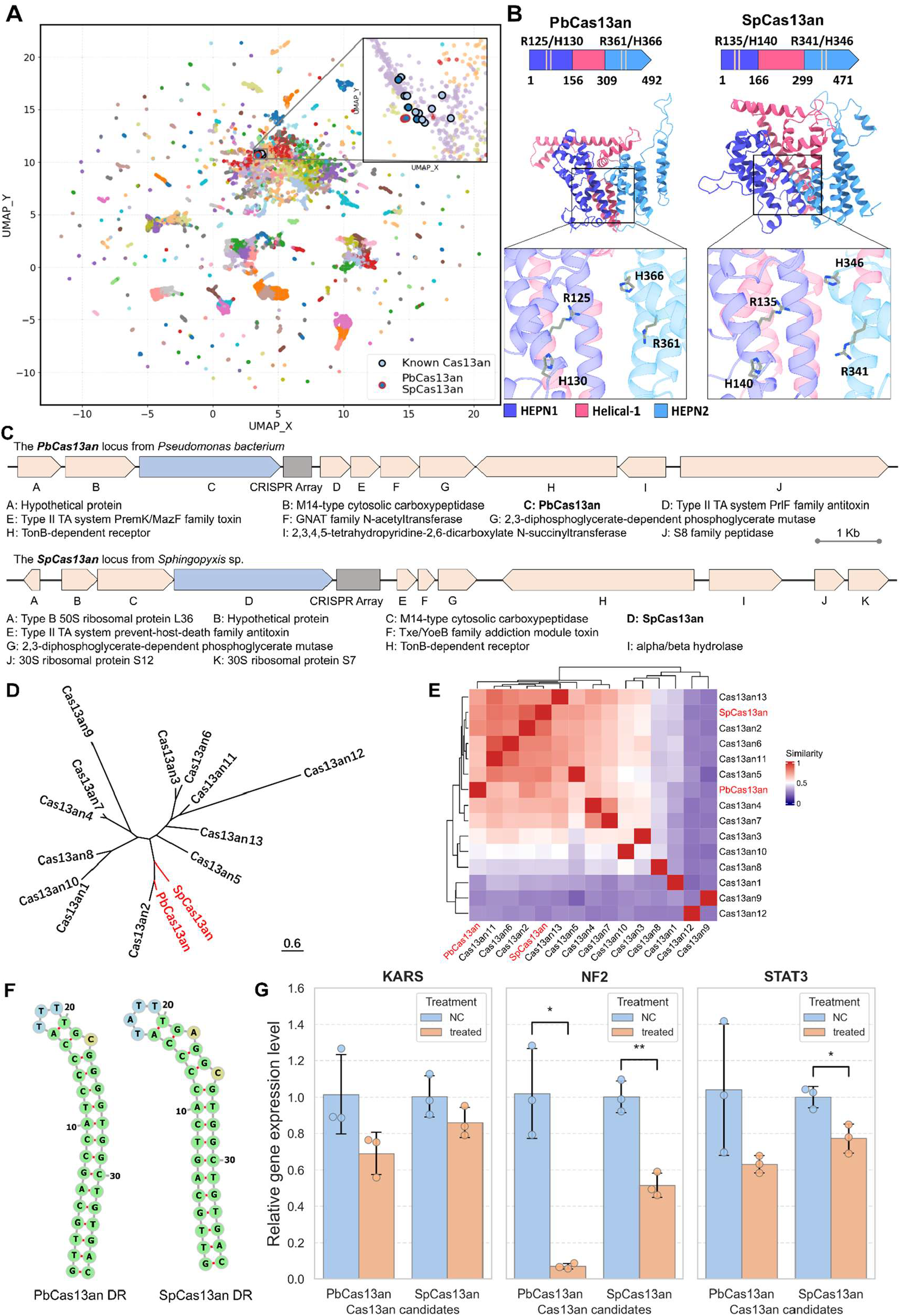
Identification and characterization of novel Cas13an candidates using sequence embedding. **(A)** UMAP dimensionality reduction of k-means clustered sequence embeddings, with zoom-in on known Cas13an-related clusters. Edges in red indicate newly identified Cas13an candidates. **(B)** Domain architectures of two novel Cas13an proteins, with annotation of the HEPN domains and the conserved RφXXXH motif. **(C)** Genomic loci of the two novel Cas13an candidates, showing neighboring genes and CRISPR array. **(D)** Maximum likelihood phylogenetic tree of the two novel Cas13an candidates alongside known Cas13an sequences. **(E)** Heatmap of TM-scores comparing structural similarity between SpCas13an, PbCas13an, and known Cas13an proteins. **(F)** Predicted secondary structures of DR sequences of PbCas13an and SpCas13an. **(G)** Quantitative analysis of RNA editing efficiencies at *KARS, NF2*, and *STAT3* loci mediated by the candidate Cas13an systems, presented as relative gene expression level.

In particular, an embedding cluster enriched with Cas13an proteins included two unreported sequences from *Sphingopyxis* sp. and *Pseudomonadota bacterium*, both of which were co-clustered with known Cas13an variants. Genomic analysis revealed adjacent CRISPR arrays near these candidate loci, supporting their role as potential RNA-targeting endonucleases. Both novel proteins also preserved critical HEPN catalytic motifs and the characteristic structural organization of Cas13 effectors. Based on these conserved biochemical and genomic features, we designated the two newly identified proteins as PbCas13an and SpCas13an.

Strikingly, the two candidate Cas13an proteins shared less than 50% sequence identity with known Cas13an effectors, yet retained both complete HEPN active sites (RφXXXH) (**Figure 4B**). Specifically, BLASTp results showed that the highest sequence identity of the PbCas13an to known Cas13an proteins was 49.9% (Cas13an2, 491 aa aligned), while the SpCas13an exhibited a maximum identity of 32.4% (Cas13an2, 471 aa aligned) (**Figure 4D**). Structural analysis further revealed that, like previously characterized Cas13an proteins, the four α-helices forming the HEPN domains in both candidates adopt a conserved N–2–3–4–1–C rearranged topology. Both candidates were also associated with orphan CRISPR arrays, further supporting their role in RNA-targeting immunity.

Furthermore, we employed ColabFold to predict the structures of PbCas13an and SpCas13an, followed by structural similarity comparisons with known Cas13an structures. The alignment revealed that both novel Cas13an candidates share notable structural similarity with the Cas13an protein family, exhibiting the relatively higher resemblance to Cas13an2 (**Figure 4E**), which is consistent with the phylogenetic analysis based on sequence data. Moreover, the DR structure of PbCas13an is identical to that of SpCas13an, both exhibiting the classic hairpin structure (**Figure 4F**). To assess the editing efficiencies of the two Cas13an candidates, we designed corresponding crRNAs targeting the *KARS, NF2*, and *STAT3* genes along with their target sequences (**Table S1**), and measured the RNA expression levels of these genes using qPCR. Real-time PCR results demonstrated that both PbCas13an and SpCas13an effectively knocked down endogenous gene expression, with particularly pronounced effects observed in *NF2*. These findings indicate that both candidates possess RNA editing activity *in vivo* (**Figure 4G**).

## Discussion

The burgeoning diversity of CRISPR-Cas systems presents both a challenge and an opportunity for understanding microbial immunity and expanding the genome editing toolbox. Traditional homology-based annotation methods, while effective for canonical systems, frequently overlook the highly divergent and non-canonical variants that are critical for unraveling the full spectrum of prokaryotic defense mechanisms. In this study, we developed BioPrinCRISPR, a class-agnostic computational framework that leverages principles of gene co-conservation, protein domain co-occurrence, and embedding-based similarity to systematically identify and characterize novel CRISPR-Cas systems across over a million bacterial genomes. Our results not only confirm the effectiveness of this principle-informed approach but also significantly broaden the known functional and architectural landscape of CRISPR-Cas effectors.

Our analysis revealed a striking architectural plasticity in CRISPR-Cas systems. The recurrent identification of fusions like the AlbA2-wHTH protein near Class I loci, and the chimeric Schlafen-Res nuclease, demonstrates that nature frequently recruits diverse domains to augment or specialize core CRISPR machinery. While domains like wHTH or Schlafen-like helicases are promiscuous and found in other cellular contexts, their consistent co-conservation with CRISPR arrays and high Cas-likeness scores provide compelling, multi-layered evidence for their functional recruitment into these defense systems. These chimeric architecture likely couples nucleic acid remodeling with target cleavage, suggesting a previously uncharacterized accessory module that may enhance canonical interference functions.

Furthermore, our embedding-based clustering approach proved exceptionally powerful for discovering novel effectors lacking discernible Pfam domains or strong homology to known Cas families. The identification of two previously unreported Cas13an-like effectors from *Sphingopyxis* sp. and *Pseudomonadota bacterium*, despite low sequence identity to known Cas13an variants, dramatically illustrates this capability. The conservation of essential HEPN catalytic motifs and characteristic structural organization logically supports their classification as true RNA-targeting Cas13an-like nucleases. These findings underscore the vast unexplored functional diversity within CRISPR-Cas systems and highlight the potential of latent features-based embeddings for de novo protein family discovery.

Collectively, our findings demonstrate that CRISPR-Cas systems exhibit far greater architectural and functional plasticity than previously appreciated. The recurrent domain fusions and novel effector architectures identified by BioPrinCRISPR represent evolutionary innovations that likely enable adaptation to diverse microbial environments and defense against a broader range of genetic threats. These discoveries contribute significantly to our understanding of the prokaryotic immune system, revealing new strategies for host-pathogen interactions and potentially uncovering novel regulatory mechanisms.

Beyond fundamental insights, the novel effectors identified by BioPrinCRISPR hold immense promise for next-generation genome engineering. The field of genome editing is continuously seeking smaller, more specific, and functionally diverse tools ^[46]^. The compact nature of some of these novel fusions, or their unique combined enzymatic activities (e.g., nuclease-helicase), could offer distinct advantages over existing CRISPR-Cas tools. For instance, a single-protein effector combining unwinding and cleavage capabilities could simplify delivery and enhance targeting specificity in complex biological systems, paving the way for applications requiring high precision and reduced off-target effects. The RNA-targeting capabilities of the novel Cas13an-like effectors also expand the utility of CRISPR technologies for RNA manipulation, a rapidly developing area in therapeutic and research applications ^[47]^.

Despite the robust computational evidence presented, it is crucial to acknowledge that in silico predictions necessitate experimental validation. Future work will focus on full biochemical and structural characterization of all identified candidates, including in vitro nuclease activity assays, target specificity profiling, and investigation of their in vivo interference capabilities against foreign nucleic acids. Detailed structural analyses of the identified fusions will also provide critical insights into their unique mechanisms of action. Moreover, applying BioPrinCRISPR to an even wider array of metagenomic datasets and under-explored microbial niches promises to uncover additional, unforeseen CRISPR-Cas diversity, further enriching our understanding of microbial immunity and expanding the biotechnological repertoire for genome engineering.

In conclusion, BioPrinCRISPR represents a powerful and validated framework for moving beyond homology-constrained discovery in the CRISPR-Cas field. By prioritizing conserved biological principles, we have unlocked a trove of novel Cas proteins and domain architectures, providing unprecedented insights into the functional evolution of microbial defense and laying a foundational roadmap for the development of innovative genome editing technologies.

## Methods

### Data Collection and Preprocessing

All publicly available prokaryotic genome assemblies were downloaded from the NCBI database (https://www.ncbi.nlm.nih.gov/) as of 1 August 2024. This dataset encompasses both complete and draft genomes across diverse prokaryotic taxa. In addition, metagenome-assembled genomes (MAGs) from unpublished datasets were included, obtained through internal projects and collaborative efforts. All genomes were subjected to standardized quality control procedures, including assessments of completeness and contamination, prior to downstream analysis.

### Computational Framework: BioPrinCRISPR

#### CRISPR Array Detection and Preprocessing

To detect CRISPR arrays across prokaryotic genomes, we employed the MinCED (version 0.4.2) ^[31]^, which is optimized for high-throughput array identification based on spacer–repeat patterns. A custom Python pipeline was developed to automate array detection across large-scale bacterial genome datasets in parallel.

For each genome assembly in our dataset (n = 1,067,859), the FASTA files were processed using MinCED with adjustable parameters for array detection: minimum number of repeats (--minNR), repeat length (--minRL, --maxRL), and spacer length (-- minSL, --maxSL). Default values were set to a minimum of 3 repeats, repeat lengths between 11–80 bp, and spacer lengths also between 11–80 bp, unless specified otherwise. These thresholds are consistent with reported biological characteristics of known CRISPR arrays. Upon completion, genome files that yielded non-empty MinCED output were retained, and corresponding CRISPR array coordinates were extracted and stored in tabular format (GFF-like). Assemblies with no detected arrays or with improperly formatted output were filtered out to reduce downstream false positives.

#### CRISPR Array-Flanking Protein-Coding Gene Extraction

To investigate the genetic context of CRISPR arrays, we extracted protein-coding genes located in their flanking regions using Prodigal ^[48]^ (version 2.6.3) with a window-based scanning approach.

For each CRISPR array detected in the previous step, we defined a genomic window of ±10,000 bp centered on the array coordinates. The sequence region within this window was extracted, and the CRISPR repeat itself was masked with ambiguous bases (N) to prevent false predictions within the repeat region.

Gene prediction within the flanking window was performed using Prodigal in metagenomic mode (-p meta), which is well-suited for incomplete or highly variable genomic contexts. The predicted protein sequences were parsed from Prodigal’s FASTA output, and associated metadata (coordinates and annotations) were extracted from the GFF output stream. For each predicted coding sequence (CDS), the absolute distance to the CRISPR repeat was computed, and CDSs were ranked based on their proximity to the array. We retained the relative position of each CDS with respect to the CRISPR array as integer labels (e.g., -3, -2, -1, [array], 1, 2, 3, …), allowing for downstream modeling of conserved gene order and synteny.

To remove redundancy in CRISPR array–flanking protein pairs and to prepare high-quality inputs for downstream co-conservation analysis, we filtered the dataset to retain only those entries with non-empty CDS lists and with CRISPR arrays satisfying user-defined repeat length and repeat copy number thresholds (default: repeat length *≥* 20 bp, repeat number *≥* 3). Each CRISPR array was converted to FASTA format and paired with the amino acid sequences of its flanking CDSs, which were similarly formatted. To reduce redundancy, we constructed a unique identifier by concatenating each array sequence with its corresponding protein sequence. Duplicates were removed based on this identifier, ensuring only unique array–protein pairs were retained. Additional filtering was applied based on genomic distance or gene order proximity to the CRISPR array, depending on user-defined mode parameters: either a base-pair distance threshold (default: ≤ 10 kbp) or a flanking ORF count threshold (default: ± 5 ORFs). The resulting arrays and proteins were exported into both full and filtered FASTA files, which were further deduplicated using SeqKit ^[49]^ (seqkit rmdup, version 2.8.2) to eliminate internal sequence redundancy.

#### Co-Conservation-Based CRISPR-Associated Protein Filtering

To prioritize CRISPR-associated proteins with evidence of functional linkage, we performed a co-conservation analysis that quantifies the co-occurrence patterns between CRISPR arrays and their neighboring proteins across genomes. This approach is based on the hypothesis that proteins consistently observed near CRISPR arrays in a statistically non-random fashion are more likely to be functionally associated with CRISPR systems.

We first constructed undirected, unweighted networks for CRISPR arrays and flanking proteins using pairwise co-clustering data derived from sequence similarity and genomic proximity. Each input cluster file (array or protein) contained pairwise links inferred from sequence-based clustering, performed using either MMseqs2 (version 15.6f452) ^[50]^ or CD-HIT (version 4.8.1) ^[51]^ depending on dataset size. The resulting links were then parsed and converted into edge lists. Using NetworkX ^[52]^ (version 3.1), we created initial graph structures where nodes represent unique sequence identifiers and edges indicate co-clustering. To eliminate noise, we removed all connected components smaller than a user-defined threshold (default: 5 nodes), and discarded self-loops to avoid trivial redundancy.

The co-conservation analysis was implemented as a combinatorial search across all pairs of CRISPR array and protein clusters. For each array–protein cluster pair, we computed the overlap in genome assemblies (i.e., whether a specific array–protein pair co-occurs in the same genomic context). A core component of this analysis is the coverage score, which quantifies the fraction of protein instances in a given cluster that co-occur with CRISPR arrays across genomes. Only protein clusters achieving a minimum co-conservation coverage threshold (default: 0.3) were retained for further analysis. The final retained protein set consisted of nodes with statistically supported co-occurrence with CRISPR arrays. We updated the original protein network by preserving only the remaining high-confidence protein nodes and their associated edges. The resulting filtered network was serialized into GraphML format for downstream visualization and domain annotation. Additionally, representative protein identifiers (i.e., central nodes in each cluster) were extracted to serve as candidates for subsequent domain-level co-occurrence profiling.

#### Domain Co-occurrence Network Construction

To investigate co-occurrence patterns of protein domains, we constructed both directed and undirected domain-level networks based on HMMER searches against the Pfam-A HMM database (version 37.1). For each query protein, domain architectures were parsed from the --domtblout output file, retaining only hits with defined coordinates and converting relevant fields to numeric types for downstream filtering.

To reduce redundancy from overlapping domain hits, a domain-merging strategy was applied: adjacent domains with more than 50% coordinate overlap (relative to either domain length) were considered redundant, and the domain with the higher conditional E-value was removed. This filtering step was repeated iteratively until no significant overlaps remained.

For the directed network, the non-redundant domains in each protein were ordered from N- to C-terminus based on start positions. Consecutive domain transitions were extracted to define ordered domain pairs, and proteins with at least one such pair were used for edge construction. Domain pairs across all proteins were aggregated to compute edge frequency and domain-specific occurrence. A directed graph was then built using NetworkX, where nodes represent Pfam domains and edges represent directional transitions between them within proteins.

In parallel, to explore domain-level functional associations independent of directionality, we constructed an undirected co-occurrence network using the same filtered HMMER annotations, focusing specifically on retained CRISPR-associated loci. Pairwise combinations of non-overlapping domains within the same protein were recorded, and only domain pairs observed in at least a user-defined number of proteins and exceeding a minimum co-occurrence threshold were retained.

The undirected network was also constructed using NetworkX, with nodes representing unique Pfam domains and edges denoting statistically supported co-occurrence events. Edge weights captured both the number of proteins sharing a given domain pair and their relative co-occurrence frequency. Additional annotations—such as domain novelty (compared to reference CRISPR-Cas datasets), functional descriptions, and protein length distributions—were integrated as node attributes. Isolated nodes and weakly supported edges were removed to simplify the final network.

#### Prioritization of Domains via Undirected Network Analysis

To identify protein domains potentially associated with CRISPR systems, we constructed an undirected co-occurrence network based on filtered HMMER annotations. Each node in the network represented a unique Pfam domain, while edges denoted domain co-occurrence within the same protein. The network was imported from a GraphML file using the NetworkX Python library, with associated node and edge attributes reflecting domain and domain-pair occurrences.

To quantify the relevance of individual domains and their associations, we assigned weights to both nodes and edges. Node weights corresponded to the number of unique proteins containing the respective domain, whereas edge weights reflected the number of proteins in which the connected domain pair co-occurred. These values were extracted directly from the GraphML attribute fields, where protein identifiers were explicitly listed for each node and edge.

To further assess the similarity of unknown domains to known Cas proteins, we applied Node2Vec (version 0.5.0) for graph-based embedding of domains into a 64-dimensional feature space. The embeddings were trained using biased random walks (walk length = 30, number of walks = 200). A subset of Pfam domains previously annotated as Cas-related—defined as occurring at least 10 times in known Cas loci—served as reference points. For each domain in the network, we computed the cosine similarity between its embedding vector and the centroid of the known Cas-domain vectors, thereby assigning a “Cas-likeness” score to each node.

To visualize structural patterns within the embedded space, we projected all domain vectors into two dimensions using t-distributed stochastic neighbor embedding (t-SNE) with a perplexity parameter of 30. In the resulting scatter plot, known Cas domains were highlighted in red, while all other domains were shown in gray. This allowed for qualitative assessment of clustering and identification of domains exhibiting Cas-like properties.

#### Embedding-Based Clustering and Unannotated Proteins

To explore functional diversity among proteins lacking Pfam annotations, we employed a two-step computational pipeline involving language model-based embeddings and unsupervised clustering. Protein sequences with no Pfam-assigned domains were first embedded using the ESM-2 protein language model (33-layer, 650M parameters; Facebook AI Research), pre-trained on UniRef50. Full-length amino acid sequences were tokenized using the accompanying ESM tokenizer and input into the model using GPU-accelerated inference. For each protein, we extracted a fixed-length representation by averaging the final-layer hidden states across all residue positions. Reference protein embeddings were prepared similarly and served as a comparative backbone to contextualize the Pfam-unannotated proteins.

The resulting embedding vectors were concatenated and L2-normalized prior to clustering. We applied the k-means algorithm (scikit-learn v1.3.0) for unsupervised partitioning of the embedding space. When the optimal number of clusters was unknown, we conducted a systematic k-means optimization over a specified range, evaluating clustering quality using both the elbow method (inertia) and average silhouette scores. Final clustering assignments were derived from the optimal k value and saved alongside sequence identifiers for further inspection.

To visualize the resulting high-dimensional protein space, we used Uniform Manifold Approximation and Projection (UMAP, n_neighbors=15, min_dist=0.1) to reduce the embedding dimensionality to two. UMAP was performed on normalized vectors, preserving local and global structure. Cluster assignments were overlaid on the 2D projection using distinct colors. Proteins of special interest (e.g., Cas13an candidates and other curated IDs) were highlighted explicitly in the scatterplot and annotated using collision-free text labeling. All plots were rendered using Matplotlib and Seaborn, and reproducible scripts were implemented to allow dynamic inspection of protein relationships based on embedding similarity.

### Protein Candidate Analysis

Protein candidate sequences were subjected to multiple sequence alignment using MAFFT ^[53]^ (version v7.520) with the L-INS-i algorithm, incorporating 1000 bootstrap replicates to assess alignment robustness. Phylogenetic reconstruction was conducted employing the maximum likelihood method implemented in IQ-TREE ^[54]^ (version v2.2.6). The optimal substitution model (Q.pfam+F+I+R7) was selected based on Bayesian Information Criterion (BIC) scores. Resulting phylogenies were further refined and annotated using the EvolView platform for enhanced graphical representation.

Tertiary structures of candidate proteins were predicted through ColabFold ^[55]^, leveraging deep learning-based homology modeling approaches. Predicted models were structurally aligned against experimentally resolved homologs using US-align ^[56]^ (version 20241108).

Sequence conservation and motif identification were visualized via sequence logos generated by WebLogo ^[57]^. For crRNA secondary structure prediction, ViennaRNA ^[58]^ package was employed, and predicted structures were graphically rendered using R2DT^[59]^ to provide standardized and interpretable RNA structural diagrams.

### Plasmid Construction, Cell Transfection, and Quantitative Analysis of Novel Cas13an Candidates

#### Plasmid Vector Construction and gRNA Design

The human codon-optimized sequences of PbCas13an and SpCas13an, each tagged with bipartite nuclear localization signals (bpNLS) at both N- and C-termini, were synthesized (GCATbio Corporation, China) and cloned into a modified PX458 vector backbone (Addgene plasmid #48138) using the ClonExpress II One Step Cloning Kit (Vazyme, C112-01). The gRNA expression cassette, driven by the U6 promoter, was engineered to include a BsaI Golden Gate cloning site for spacer insertion. Target sites within *KRAS, NF2*, and *STAT3* transcripts (listed in Supplementary Table S1) were selected based on predicted RNA accessibility and specificity. Complementary oligonucleotides encoding spacer sequences were annealed and inserted into the gRNA cassette via *Bsa*I digestion followed by T4 DNA ligase-mediated ligation. The direct repeat sequence was positioned at the 3′ end of the spacer in the gRNA cassette.

#### Cell Culture, Transfection, and Gene Expression Detection

HEK293T cells were cultured in high-glucose Dulbecco’s Modified Eagle Medium (DMEM) supplemented with sodium pyruvate (Gibco, 11995065), 10% fetal bovine serum (Thermo Fisher Scientific, 10100147), and 1× penicillin-streptomycin (Thermo Fisher Scientific, 15140148) under standard conditions (37°C, 5% CO_2). Cells were initially expanded in 100 mm dishes and subsequently seeded into 12-well plates at a density of 1×10^5 cells per well one day prior to transfection. When cells reached approximately 70% confluency, plasmid DNA encoding PbCas13an or SpCas13an and their corresponding gRNAs were transfected using Lipo8000 reagent (Beyotime Biotechnology, C0533) according to the manufacturer’s instructions. Seventy-two hours post-transfection, cells were harvested for RNA extraction.

Total RNA was isolated using the RNAsimple Total RNA Kit (TIANGEN Biotech, 4992858), following the manufacturer’s protocol. Complementary DNA (cDNA) synthesis was conducted using HiScript II Q Select RT SuperMix for qPCR (+ gDNA wiper) (Vazyme, R233-01). Target gene expression was quantified by real-time PCR using AceQ qPCR SYBR Green Master Mix (Vazyme, Q111-02). Relative transcript levels were normalized to β-actin and analyzed using the 2^-ΔΔCt method^[60]^. Primer sequences for quantitative PCR are listed in Supplementary Table S2.

## Data and Code Availability

All source code for the BioPrinCRISPR pipeline developed in this study is publicly available on GitHub at https://github.com/Bio-bbhe/BioPrinCRISPR. The interactive web application is available at https://crispr-384817688195.asia-east2.run.app. The publicly available prokaryotic genomes used in this analysis were downloaded from the NCBI database; a complete list of accession numbers is provided in Supplementary Table S3.

## Acknowledgments

We acknowledge the BGI Research’s High-Performance Computing Center for providing computational resources and technical support.

## Author contributions

B.B.H., C.Q. and H.X.L. conceived the study. B.B.H. and C.Q. curated data, developed algorithm, analyzed the data, and prioritized the candidates. D.L., B.B.H. and C.Q. contributed to the web development. Y.Y.F. and F.R.W. conducted experimental validation. B.B.H., C.Q., H.X.L. and Y.Y.F. wrote the original draft of the manuscript. Z.A.W. contributed to the draft revision. D.W., Y.X.L., Y.Z, Yong Zhang, H.X.L. and Z.Y. supervised the study. All authors reviewed and approved the final manuscript.

